# Ion channel distributions in cortical neurons are optimized for energy-efficient active dendritic computations

**DOI:** 10.1101/2021.12.11.472235

**Authors:** Arco Bast, Marcel Oberlaender

## Abstract

The mammalian brain has an enormous demand for energy, which is thought to impose strong selective pressure by which the neurons evolve in ways that ensure robust function at minimal energy cost. However, which principles drive the ion channel distributions in the dendrites to implement different neuronal functions is yet unclear. Here we found that an energy-efficient generation of dendritic calcium action potentials in cortical pyramidal neurons requires a low expression of slow inactivating potassium channels. We demonstrate that this relationship between energy cost and neuronal function is independent of the dendritic morphology and the expression patterns of other ion channels that implement additional perisomatic and dendritic functions. Moreover, we found that calcium action potentials can arise from a wide spectrum of ion channel expression patterns, including configurations with high potassium channel densities in the dendrites. These configurations can account equally well for the characteristic intrinsic physiology of the pyramidal neurons. However, only configurations with low potassium channel densities in the distal dendrites are observed empirically. Thus, our findings indicate that cortical neurons do not utilize all theoretically possible ways to implement their functions, but instead select those optimized for energy-efficient active dendritic computations.

## Introduction

It is thought that neurons in the cerebral cortex evolved to be energy-efficient in neural computation^1, 2^. Most of the cortical energy costs result from the generation and transmission of electrical and chemical signals within and between the neurons^3^. For example, the initiation of action potentials (APs) in the axon is metabolically expensive, consuming more than 20% of the energy in the gray matter of the rodent brain^4, 5^. Experimental studies^6, 7^ and simulations of single compartment models^8^ found that the energy costs for AP generation in mammalian neurons are remarkably close to the theoretically predicted minimum^9^, whereas the energy consumption of APs in invertebrates is far from being optimal^8, 10^. These observations indicate that biophysical parameters in the axon of cortical neurons – e.g. the kinetics of ion channels^11^ or the expression patterns of sodium (Na^+^) and potassium (K^+^) channels^12^ – are optimized for generating APs with minimal energy costs. However, whether optimization of biophysical parameters could represent a general principle by which cortical neurons implement energy-efficient signal processing in the dendrites is unknown.

One major challenge for answering this question is that cortical neurons, and particularly the pyramidal neurons in layer 5 (L5), express a variety of voltage-gated ion channels in specific dendritic domains^13, 14^. As a result, the pyramidal neurons can perform active computations in their dendrites, for example by generating calcium (Ca^2+^) APs around the main branch point of the apical dendrite^15–18^. Distal dendritic inputs, which normally appear greatly attenuated at the cell body (i.e., soma), must cross a high threshold to evoke calcium APs^17^, but can then generate bursts of axonal APs^19^. Thereby, L5 pyramidal neurons can act as cellular ‘coincidence detectors’ by switching their axonal output from single AP to burst firing when proximal and distal inputs arrive synchronously^19^. Thus, while single compartment models are sufficient to assess whether biophysical parameters are optimized in the axon for energy-efficient generation of single APs^8^, deriving such information for signal processing in entire pyramidal neurons requires models that account for both the particular morphology of the dendrites and the respective expression patterns of the various ion channels along them.

Previous models of L5 pyramidal neurons were able to capture certain aspects of the complex intrinsic physiology separately^20–22^. However, generating models that jointly capture the dendritic, perisomatic and axonal physiology remains challenging. To the best of our knowledge, a model accomplishing this task is currently only available for one dendritic morphology reconstructed from an *in vitro* labeled L5 pyramidal neuron^13^. The ion channel distributions derived for this morphology failed however to reproduce the intrinsic physiology observed for other L5 pyramidal neurons^23^. Therefore, it is thought that L5 pyramidal neurons – and neurons in general – establish their proper function by compensating morphological variability with variability in ion channel expression patterns^24^. However, conclusive answers to the questions – *Which principles govern the relationships between morphology, channel expression and neuronal function, and is energy efficiency one of these principles?* – remain unknown.

Here we quantitatively address these questions for the major output cell type of the cerebral cortex^25^, the subcortically projecting layer 5 pyramidal tract neurons (L5PTs). For this purpose, we generate biophysically detailed multi-compartmental models for a set of *in vivo* labeled L5PTs^26–28^ that is representative for the morphological variability of this class of neurons in the vibrissa-related part of the rat primary somatosensory cortex **(Fig. S1)** – the barrel cortex. We demonstrate that we can generate hundreds of thousands of models with very different biophysical parameters for each of the morphologies, and that all of these models capture equally well the dendritic, perisomatic and axonal physiology that is observed empirically for the class of L5PTs in the rat barrel cortex^13^. This comprehensive set of biophysically, morphologically and physiologically realistic models allows us assess the energy costs for characteristic functions of the L5PTs, and to predict how ion channels would be expressed along the dendrites if they were optimized for energy-efficient signal processing.

## Results

We generated multi-compartmental models for *in vivo* labeled dendrite morphologies **(Fig. 1a)** to capture 40 features that characterize the complex intrinsic physiology of L5PTs^13^ **(Table S1)**. These features reflect properties of APs in response to brief and prolonged somatic current injections (i.e., perisomatic physiology), as well as backpropagation of APs into the apical dendrite, dendritic calcium APs, and bursts of APs in response to coincident somatic and dendritic current injections (i.e., dendritic physiology) **(Fig. 1b)**. Each model comprises 32 biophysical parameters **(Fig. 1c)** to describe passive electrical properties, intracellular calcium buffer kinetics, and the respective distributions and densities of a diverse set of ion channels in the soma, axon, basal and apical dendrites **(Table S2)**. These biophysical parameters and physiological features were reported by Hay et al.,^13^, representing so far the only modelling attempt that could successfully capture the perisomatic and dendritic physiology of a L5PT. Here, we modified the genetic algorithm that Hay et al.,^13^ developed for parameter fitting to be suitable for exploring the biophysical parameter space (Methods). This modification allowed us to identify parameter values that yield responses to the somatic and dendritic current injections that are consistent with those observed empirically across L5PTs. We consider models that are consistent in all 40 features as ‘acceptable’ **(Fig. S1c)**. However, even extensive exploration of the parameter space generally failed to generate acceptable models for any of the L5PT morphologies that we tested, particularly for capturing the features of calcium APs and bursts of APs during coincidence detection.

**Fig. 1:**
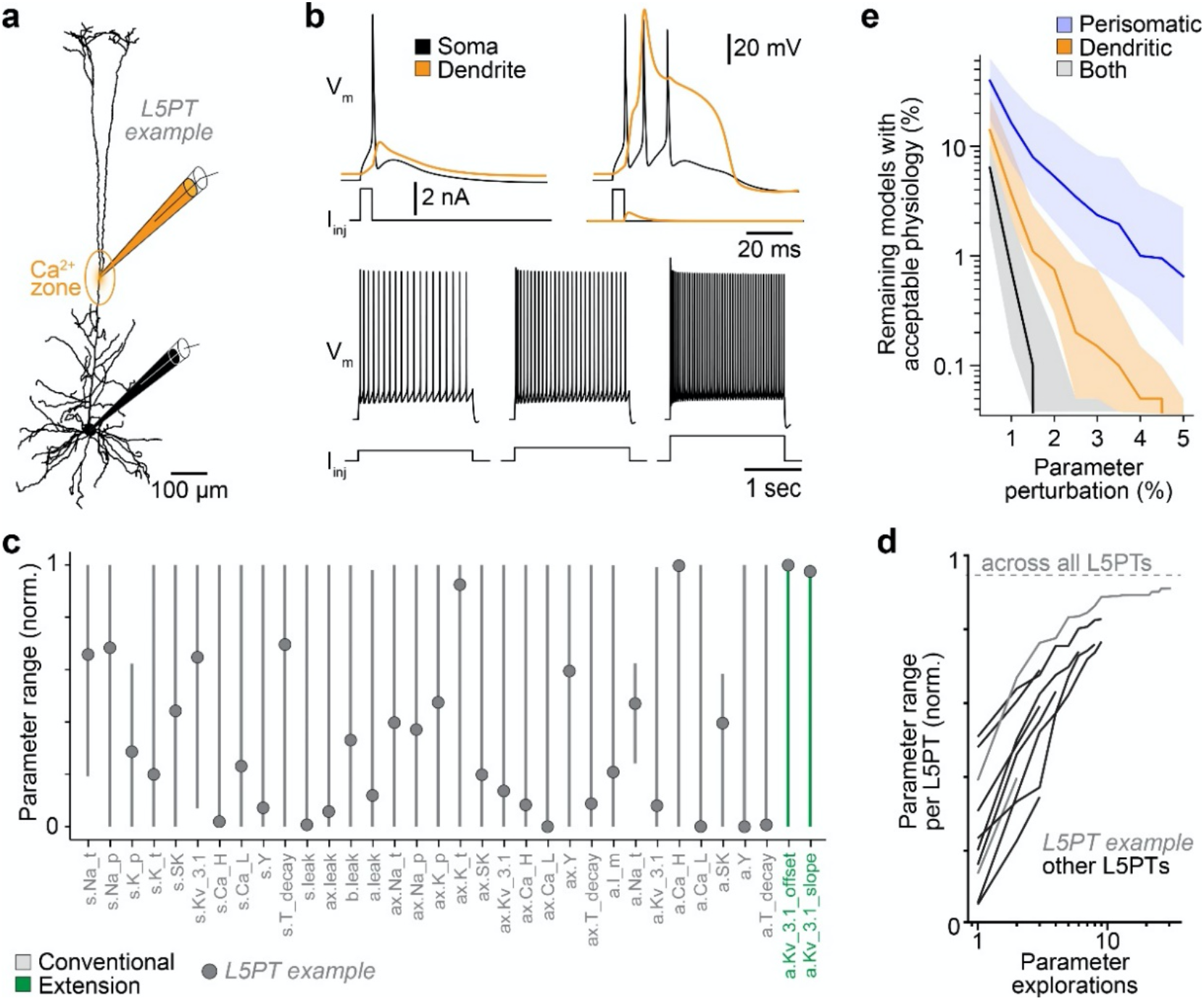
Generating functionally realistic models for *in vivo* labeled cortical neurons. **a.** Example dendrite morphology from the 12 layer 5 pyramidal tract neurons (L5PTs) for which we generated multi-compartmental models **(Fig. S1a)**. The main branch point of the apical dendrite reflects the initiation zone for calcium (Ca^2+^) action potentials (APs). **b.** Example simulations of current injections into the soma (black) and/or Ca^2+^ zone (orange) of a model for the L5PT from panel a. Consistent with empirical recording data **(Table S1)**, all models analyzed here support the characteristic dendritic (top) and perisomatic (bottom) physiology of L5PTs **(Fig. S1c)**, including backpropagation of APs into the apical dendrite (top-left), calcium APs and burst firing when inputs to the soma and Ca^2+^ zone coincide (top-right), and regular AP firing of increasing frequencies in response to sustained current injections of increasing amplitude (bottom). **c.** Range of biophysical parameters across all models with acceptable dendritic and perisomatic physiology (n=4.2 millions) representing the passive leak conductance and the density of Hodgkin-Huxley type ion channels on the soma (s), basal dendrite (b), apical dendrite (a), and axon initial segment (ax). We extended the 32 parameters proposed by Hay et al.,^13^ by two biophysical parameters (green) that allows the density of the fast non-inactivating potassium channel (Kv3.1) to decay along the apical dendrite (slope) until a minimum is reached (offset)^30^. Parameters are normalized to their biophysically plausible range (Methods). Dots represent the example model from panel b. **d.** Each exploration of the parameter space leads to new acceptable models that extend the parameter range for each morphology. Across all morphologies (dashed line), 95% of the plausible parameter ranges are covered (i.e., mean range across all parameters across all models). The parameter ranges covered by individual morphologies approach that across all morphologies with increasing numbers of parameter space explorations. **e.** Perturbing the parameters of acceptable models results in physiology that is no longer within the empirically observed range. Dendritic physiology is more sensitive to perturbations than perisomatic physiology. ‘Remaining models’ refers to the fraction of models that were acceptable out of randomly sampled models from the parameter space where each parameter was perturbed by ≤1%, 2%, etc. Shaded are is 25^th^ to 75^th^ percentile, bold line is median.

We therefore considered that some important biophysical parameters that are necessary to capture the dendritic physiology of L5PTs might be missing in the models. Two previous studies reported that the expression of slow inactivating potassium channels steeply decays with soma distance along the apical dendrite of L5PTs^29, 30^. The functional relevance of this observation remained however unclear to this date. Remarkably, when we added two parameters to incorporate the slope and offset of such potassium channels (Methods), exploration of the hence extended parameter space resulted in models with acceptable dendritic physiology **(Fig. 1a-c)**. In fact, additional explorations continued to produce new sets of models with unique combinations of biophysical parameter values – i.e., with different ion channel distributions and densities, and passive electrical properties. Thus, as this extended biophysical parameter space increases the likelihood to generate acceptable models, a decay of slow inactivating potassium channels along the apical dendrite appears to be key for the robust function of L5PTs, particularly for the generation of calcium APs and bursts of APs during coincidence detection.

We explored the extended parameter space first for values that yield models with acceptable dendritic physiology – i.e., the responses of each model to two different current injections (top row in **Fig. 1b**) are validated against the ten features that characterize backpropagating APs, calcium APs and bursts of APs **(Fig. S2)**. Exploration of the parameter space from a new seed point generally identified additional acceptable models. We therefore continued the exploration until the range of parameter values reached a plateau **(Fig. 1d)**. For the 12 morphologies used in this study, we thereby generated more than 11 million models with acceptable dendritic physiology that covered 98% of the plausible parameter ranges. Thus, surprisingly, only very few of the biophysical parameters were strongly constrained by the dendritic physiology. Instead, most parameters in the soma, dendrites and axon ranged from the minimal plausible value to the maximal one. We therefore explored parameter values that yield acceptable models for both the dendritic and perisomatic physiology. For this purpose, we validated each model against all of the 40 features that characterize the intrinsic physiology of the L5PTs^13^ **(Table S1)**. We again continued the exploration until the parameter ranges reached a plateau, which resulted in more than 4.2 million acceptable models that still covered 95% of the plausible ranges **(Fig. 1c)**. Thus, even a 4-fold increase of the functional constraints only minimally narrowed down the volume of the parameter space that comprises acceptable models.

Does the wide range imply that the biophysical parameters in L5PTs are only weakly constrained by their intrinsic physiology, or could it simply reflect morphological differences? To address this possibility, we computed the parameter range across all models for individual morphologies. Even for the same morphology, the parameters covered up to 92% of the plausible range **(Fig. 1d)**, thereby rivaling the parameter range observed across all morphologies combined. Does this mean that L5PTs are insensitive to changes in the distributions and densities of particular ion channels? To address this possibility, we perturbed the biophysical parameters of acceptable models. Perturbing the parameters by 1% substantially changed the resulting intrinsic physiology, so that more than 99% of the models were no longer acceptable **(Fig. 1e)**. Thus, while the spectrum of channel expression patterns by which L5PTs could implement their characteristic intrinsic physiology is equally widespread for the different morphologies, the respective implementations within this spectrum are specific for each morphology.

Next, we investigated how L5PTs dynamically utilize ion channels – i.e., how their opening and/or closing contributes to the currents flowing in and out of the soma, axon and dendrites. For this purpose, we focused on the generation of calcium APs during coincidence detection, as this mechanism requires proper channel expression patterns that capture both the perisomatic and dendritic physiology. For each of the acceptable models, we quantified the respective charge exchanged through the ion channels that are expressed at the initiation zone for calcium APs – i.e., around the main branch point of the apical dendrite **(Fig. 2a)**. Although this was not an objective of the genetic algorithm, all models first used sodium and then calcium channels for depolarization, giving rise to a Na^+^/Ca^2+^-complex as observed empirically^31^. Surprisingly, each model utilized the sodium, calcium and potassium channels very differently to generate calcium APs, even though the resulting shapes of the calcium APs and axonal AP bursts were very similar **(Fig. 2a)**, and well within the experimentally observed range **(Table S1)**. For example, some models relied entirely on high-voltage activated calcium channels (Ca_HVA) for depolarization^32^, and used a superposition of calcium-dependent (SK) and fast noninactivating potassium channels (Kv3.1) for hyperpolarization. Other models relied on low-voltage activated calcium channels (Ca_LVA) for depolarization, while using either the muscarinic (M) or Kv3.1 potassium channel for hyperpolarization. These observations generalized across morphologies **(Fig. 2b)**. Thus, largely disparate channels utilizations can implement the same functions in L5PTs.

**Fig. 2:**
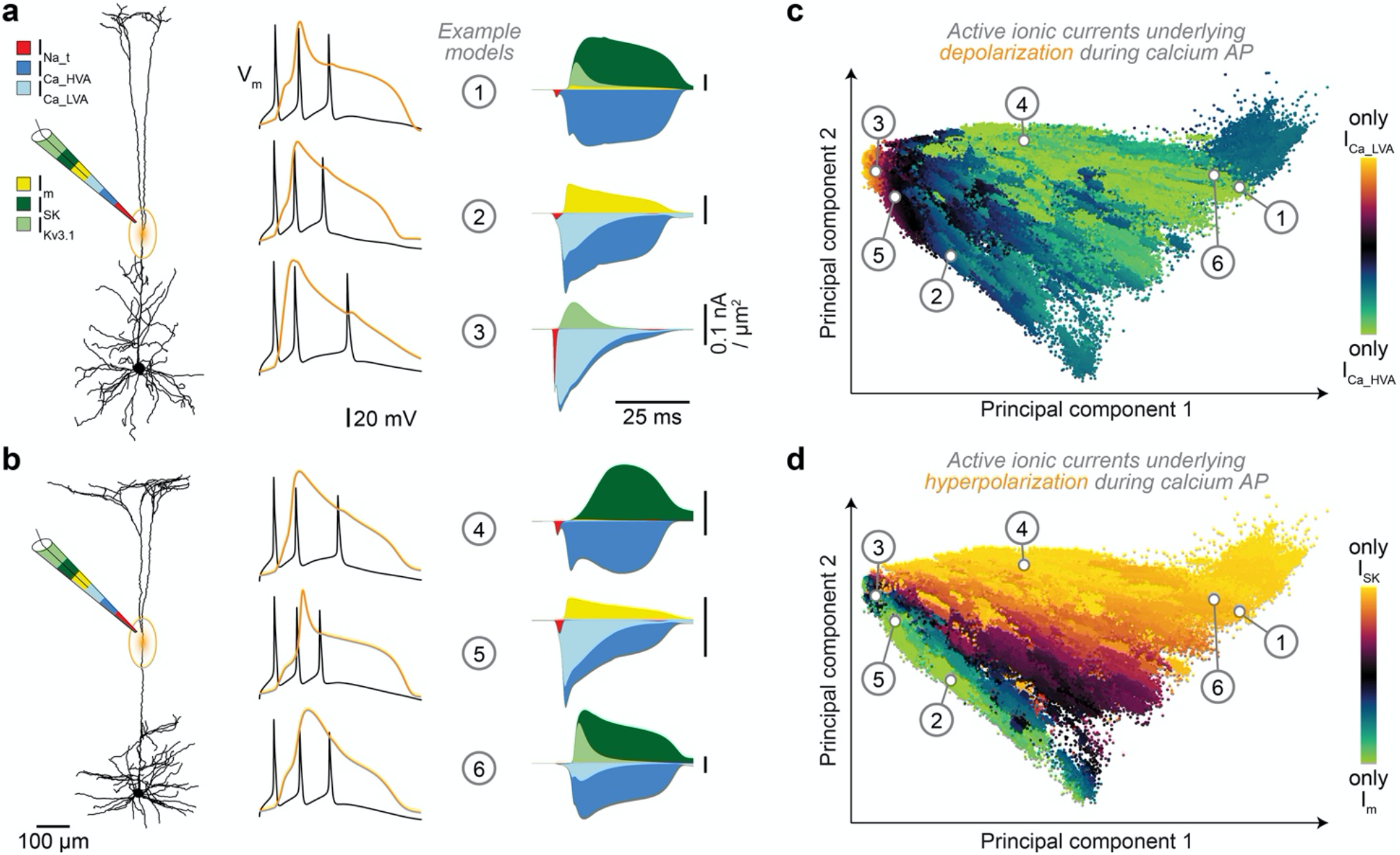
Active computations in cortical dendrites arise from a wide spectrum of ionic currents. **a.** Three example models of the same morphology (left) that generate Ca^2+^ APs and axonal bursts of APs during coincidence detection within the experimentally observed range (middle), but which utilize very different superpositions of ion channels to achieve this function (right). Hyperpolarizing currents are plotted upwards (i.e., calcium-dependent (SK), fast non-inactivating (Kv3.1), and muscarinic (M) potassium channels), depolarizing currents downwards (i.e., low- and high-voltage activated calcium channels (Ca_LVA and Ca_HVA) and sodium (Na_t) channels). **b**: Example models with diverse channel unitizations similar to those in panel a, but for a different L5PT morphology. **c**. Principal component space (Methods) illustrating the wide spectrum of active ionic currents that can theoretically underlie dendritic calcium APs, color-coded by the contribution of the two most strongly utilized depolarizing channels. Visualized are 15.5 million models that capture the characteristic dendritic physiology of L5PTs, and which include the 4.2 million models that additionally capture the characteristic perisomatic physiology **(Fig. S3)**. **d.** Same as in panel c, but color-coded for the two channels with the strongest contribution to hyperpolarization.

To analyze channel utilization more systematically, we represented each of the millions of models in a six-dimensional space, where each dimension reflects the charge exchanged through one of the channels that contribute to depolarization or hyperpolarization during calcium APs (Methods). The first two principal components of this space captured more than 90% of the variability in channel utilization. While the utilization scenarios described above were located at the boundaries of the principle component space **(Fig. 2c)**, the entire space was densely filled with models from the different morphologies. This indicates that the example scenarios reflect rather extreme cases of a wide spectrum by which L5PTs can utilize superpositions of different ion channels to implement the same function **(Fig. 2d)**. Therefore, we considered that the wide spectrum of channel utilization could reflect the wide spectrum of channel expression that we observed across models. However, the more a particular channel was expressed did not necessarily imply that it was utilized more **(Table 1)**. For example, the expression of low-voltage activated calcium channels might be much higher than that of the high-voltage activated ones, but the latter could still contribute the most to the depolarizing currents during the initiation of calcium APs. Similarly, models that expressed a particular channel more than others did not necessarily rely on it more to generate calcium APs. Thus, channel expression does not directly translate into how L5PTs utilize channels to implement their functions.

**Table 1:** Weak link between ion channel expression and utilization. We define ion channel utilization as the fraction of the charge exchanged through the specific channel in comparison to the charge exchanged through all other depolarizing or hyperpolarizing channels during a dendritic calcium AP that is generated around the main branch point (BP) of the apical dendrite. The relationship between ion channel expression and channel utilization is complex. First, comparing the expression of two ion channels in the same model illustrates that the channel that is expressed more is not necessarily utilized more. For example, model 2 expresses the low-voltage activated calcium (Ca_LVA) channel with a maximum conductance 54 times higher than that of the high-voltage activated calcium (Ca_HVA) channel, but the latter contributes the majority (69%) of the depolarizing currents during the Ca^2+^ AP. Second, comparing the expression of the same channel between two different models illustrates that the model that expresses a particular channel more does not necessarily rely more on it to perform its function. For example, model 1 expresses four times more of the fast non-inactivating potassium (Kv3.1) channel compared to model 3. However, the Kv3.1channel contributes 11% to hyperpolarization in model 1 versus 99% in model 3.

| | channel expression @ BP<br>max conductance (nA/μm) | | | channel utilization @ BP<br>current contribution (%) | | |
| --- | --- | --- | --- | --- | --- | --- |
|  | model 1 | model 2 | model 3 | model 1 | model 2 | model 3 |
| **depolarization** |  |  |  |  |  |  |
| Ca_HVA | 49.83 | **35.03** | 3.22 | 99.12 | **69.69** | 19.92 |
| Ca_LVA | 1.49 | **1885.33** | 834.80 | 0.03 | **29.32** | 74.97 |
| Na_t | 188.06 | 165.44 | 200.17 | 0.85 | 0.99 | 5.11 |
| **hyperpolarization** |  |  |  |  |  |  |
| SK | 39.56 | 0.00 | 0.02 | 83.75 | 0.01 | 0.87 |
| M | 2.09 | 8.77 | 0.00 | 5.49 | 99.98 | 0.01 |
| Kv3.1 | **32.08** | 0.00 | **8.34** | **10.76** | 0.01 | **99.12** |

If neither diversity in morphology nor channel expression provide strong constraints for how L5PTs could implement their functions, which organizing principles underlie the wide spectrum of channel utilization? While all of the models in the spectrum can account equally well for the empirically observed dendritic physiology of L5PTs **(Fig. S3)**, we noticed large differences in the current amplitudes that the different models required to generate calcium APs. Therefore, we computed the energy cost for the generation of calcium APs across all models (Methods). Energy cost correlated strongly (Pearson R=0.94, p<0.0001, two-sided p-test) with the first principle component of the channel utilization space **(Fig. 3a)**. This relationship was independent of the dendritic morphology and whether models captured also the perisomatic physiology or not **(Fig. S4)**. Could the different energy costs reflect a simple upscaling of channel activities? If depolarizing and hyperpolarizing currents were scaled by a similar factor, the utilization of channels is essentially the same, while energy costs would differ. However, the principal component space reflects gradual changes in the relative contribution of different depolarizing and hyperpolarizing currents. For example, models where SK channels dominate hyperpolarization gradually transition along the second principle component into models dominated by M channels, while all of these models consume similar energy to generate calcium APs. Thus, irrespective of energy costs, L5PTs could implement their functions with a wide spectrum of channel utilizations, which cannot be explained by similar scaling of channel activities.

**Fig. 3:**
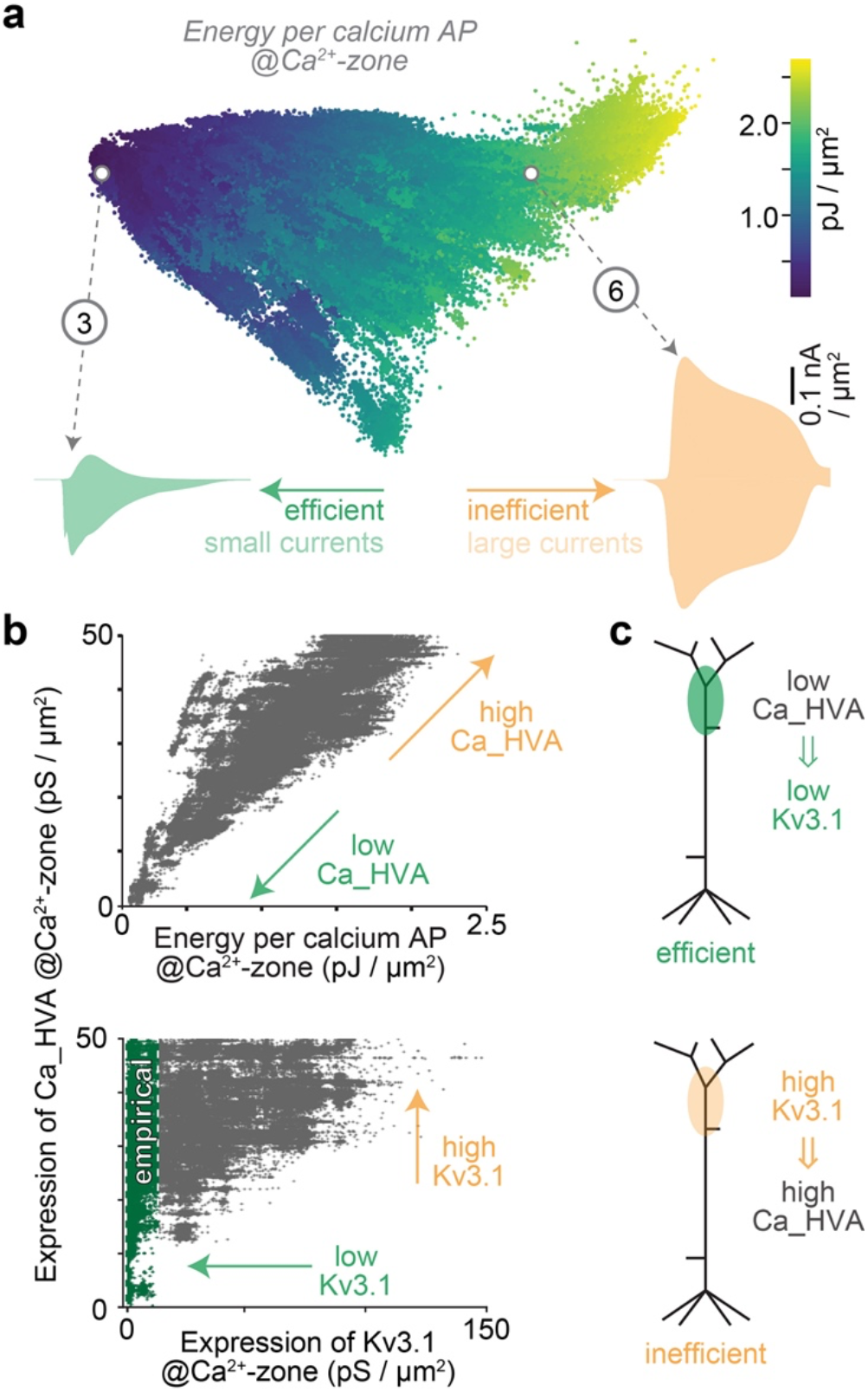
Energy-efficient signal processing constrains ion channel expression in cortical dendrites. **a.** Same principal component space as in **Fig. 2c-d**, but color-coded for the energy cost (Methods) of generating Ca^2+^ APs. Energy cost increases 10-fold along the first principal component of the theoretically possible channel utilization space. **b.** Top: Energy cost for calcium APs strongly correlates (R=0.94, p<0.0001, twosided p-test) with the expression of Ca_HVA channels in the distal apical dendrite (i.e., at the Ca^2+^ zone around the main branch point). Bottom: The expression of Ca_HVA channels restricts the expression of Kv3.1 channels. The empirically observed expression for slow inactivating potassium channels^30^, which include Kv3.1 channels, is highlighted in green. **c.** Panel b reveals that energy-efficient models generate Ca^2+^ APs with low expression of Ca_HVA, that a low expression of Kv3.1 channels is a prerequisite for a low expression of Ca_HVA channels, and hence that efficient models require a low expression of Kv3.1 channels. Thus, because models with high expression of Kv3.1 channels in the distal dendrites are inconsistent with the empirical data^30^, our findings indicate that L5PTs do not utilize these theoretically possible, but energy inefficient implementations of dendritic Ca^2+^ APs.

Given that even energy-efficient implementations of calcium APs can originate from a wide spectrum of channel expression and utilization, how will it be possible to test empirically whether the biophysical parameters of L5PTs are optimized for energy-efficient signal processing? For this purpose, we investigated whether particular expression patterns of ion channels are a prerequisite for low energy cost. We found that energy cost correlated strongly (R=0.94, p<0.0001, two-sided p-test) with the expression of high-voltage activated calcium channels across all models **(Fig. 3b)**. The higher the density of this channel at the initiation zone of calcium APs, the higher the energy cost. Surprisingly, the expression of these calcium channels restricts the expression of the fast non-inactivating potassium channels **(Fig. 3b)**. Specifically, energy-efficient models need low expression of both high-voltage activated calcium and fast non-inactivating potassium channels. Vice versa, if the expression of fast non-inactivating potassium channels is high, L5PTs need a high expression of the calcium channels to generate calcium APs, which results in higher energy cost. Hence, if energy were to constrain the implementation of signal processing in L5PTs, the density of fast non-inactivating potassium channels needs to be low in the distal apical dendrite **(Fig. 3c)**. This prediction is consistent with the observations from the studies that we used to extend the biophysical parameter space^29, 30^. Thus, the steep decay of slow inactivating potassium channels does not only ensure robust function of L5PTs by facilitating coincidence detection, a low expression of such channels in the distal apical dendrites is also the prerequisite for an energy-efficient implementation of the underlying calcium AP.

## Discussion

In this study, we investigated which principles can drive the distribution of ion channels in cortical dendrites. We found that a low expression of slow inactivating potassium channels in the distal apical dendrite is required for an energy-efficient generation of dendritic calcium APs in L5PTs. In contrast, neither neuronal function (i.e., as characterized by 40 features of the perisomatic and dendritic physiology) nor morphological properties of the dendrites require any channel density to be regulated to any specific value. Thus, each neuron has an enormous degree of freedom to implement its functions with a wide range of different ion channel distributions. It is this degree of freedom which allows cortical neurons to optimize for energy efficiency without compromising their function.

We systematically explored the ion channel expressions patterns that can give rise to the characteristic dendritic and perisomatic physiology of the L5PTs. Consistent with previous studies^13, 33^, we found that parameter values, which result in acceptable models for one morphology, can generally not be transferred to any other morphology. We also confirm that ion channel expression patterns need to be highly specific, as small perturbations lead to loss of function. However, we demonstrate that the spectrum of such specific expression patterns is widespread, such that neither morphology nor function drive the distributions and densities of any ion channel to any particular value. Contrary to conventional views^33^, morphological variability is hence not the primary source of variability in ion channel expression. This principle generalize to all scales investigated here – morphology, channel expression, channel utilization, and function. Thus, variability in one scale can only account for a minority of the variability in the other scales.

Previous work had already demonstrated that similar neuronal function can arise from largely disparate implementations. For example, experimental^34, 35^ and computational^35–37^ studies of the pyloric rhythm in the stomatogastric nervous system of cancer borealis found that different synaptic, biophysical and network parameters can give rise to virtually indistinguishable activity patterns. Furthermore, multicompartmental models of the same Purkinje cell morphology reproduced experimental activity data despite different channel expression patterns^38^. Here, we demonstrate that also for active mechanisms in the dendrites of cortical pyramidal neurons, biophysical properties do not need to be regulated to specific values to allow for proper function. This degree of freedom enables neurons to incorporate overarching principles – like energy efficiency – as determinants of ion channel distributions. Specifically, a high expression of slow inactivating potassium channels and energy-efficient dendritic calcium APs are mutually exclusive. Not a single model that expresses a high amount of this channel is energy-efficient. While we demonstrate that realistic neuronal function can arise from both high and low expression of potassium channels in the distal dendrites, energy-efficient dendritic calcium APs cannot, and strictly require a low expression. Empirical observations show that the expression of such channels falls exponentially with soma distance in L5PTs^29, 30^, and indeed reaches low values at the distal dendrites^30^, which match those predicted here for energy-efficient implementations of calcium APs. Thus, we propose that cortical neurons do not utilize all possible ways to implement their functions, but instead select those optimized for energy-efficient active dendritic computations.

Similar to the studies of single compartment models, which indicated that ion channels are optimized for energy-efficient generation of sodium APs in the axon^6–8^, our ability to generate a comprehensive set of functionally realistic multi-compartmental models indicates that such optimization generalizes to the calcium APs in the dendrites of cortical neurons. Recent studies linked dendritic calcium APs in L5PTs to perception and consciousness^39–42^. Thus, our findings might reflect a strategy that allows the cerebral cortex to evolve towards energy-efficient implementations of higher brain functions.

## Materials and Methods

### Data availability statement

Representative models for each morphology, including a documentation of all parameters and the analysis routines can be obtained from ModelDB. (https://senselab.med.yale.edu/ModelDB/; accession number: 267217).

### Multi-compartmental models

We generated and analyzed 15.5 million multi-compartmental models for dendritic morphologies that represent 12 L5PTs from the rat barrel cortex **(Fig. S1a)**. The morphologies originated from studies^26, 28, 43^ in which we had labeled individual neurons *in vivo* with Biocytin via cell-attached recordings (Wistar rats, postnatal day 25-46, male/female, Charles River). In these studies, we reported dendrite reconstructions for 91 excitatory neurons whose somata were located in the infragranular layers 5/6. Based on soma depth and 21 morphological features^27^, these neurons were objectively identified as L5PTs (n=44), or as one of the other major excitatory cell types in layers 5/6 (n=47) – i.e., intratelencephalic (L5IT), corticocortical (L6CC) or corticothalamic (L6CT) neurons^25^. From these data, we selected 12 L5PTs that were representative for the range of soma depth locations and dendritic features (e.g. path length, branch points, distance between the soma and the main branch point of the apical dendrite) observed across all reconstructed neurons of this cell type **(Fig. S1b)**. The soma-dendrite reconstructions of the 12 L5PTs were converted into multi-compartmental models. First, a simplified axon morphology was attached to the soma as described previously^23^. The axon consisted of an axon hillock with a diameter tapering from 3 μm to 1.75 μm over a length of 20 μm, an axon initial segment (AIS) of 30 μm length and diameter tapering from 1.75 μm to 1 μm diameter, and 1 mm of myelinated axon (diameter of 1 μm). Second, spatial discretization of the dendrite morphology (i.e., compartmentalization) was performed by computing the electrotonic length constant of each dendrite branch at a frequency of 100 Hz and setting the length of individual compartments in this branch to 10% of this length constant. Third, we incorporated the dynamical models of ion channels and intracellular calcium dynamics developed by Hay et al.,^13^ and assigned them to the soma, basal dendrites, apical dendrite and/or AIS, as summarized in **Table S2**. The resulting biophysical parameters were the passive leak conductance, the maximum conductance of each ion channel, and two parameters characterizing the calcium dynamics in the soma, axon, basal and apical dendrites. In addition to these free parameters, the models include the non-specific cation current Ih, which increases exponentially with soma distance to match the empirically observed voltage slope along the dendritic tree^13^. Calcium channels on the apical dendrite were expressed in high density for ±100 μm around the main branch point. Outside the hence defined calcium zone, and for oblique dendrites in general, calcium channels along the apical dendrite were expressed with 1/100^th^ and 1/10^th^ of the calcium zone density for Ca_LVA and Ca_HVA, respectively. To account for the empirically observed decay along the apical dendrite of slow inactivating potassium channels^30^, we defined a *slope* parameter for the linear decrease of the density of fast non-inactivating potassium (Kv3.1) channels with soma distance, and an *offset* parameter for the minimum density of Kv3.1 channels throughout the apical dendrite. Kv3.1 channel density on the apical dendrite is hence defined as follows:

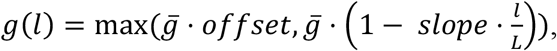

where *l* denotes the path length to the soma, *g*(*l*) the maximum conductance of the Kv3.1 channel on the apical dendrite as a function of path length to the soma, 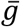 the maximum conductance of the Kv3.1 channel on the apical dendrite closest to the soma, and *L* the maximum path length (i.e., the distance of the dendritic segment with the largest distance to the soma). Furthermore, we added a parameter to scale the diameters of apical dendrites, which were reconstructed at the resolution limit of light microscopy. The resulting range for the diameters of the apical dendrite (ranging from 1.3 to 4.2 at the main branch point) was consistent with empirical measurements in rat somatosensory^44^ and visual^45^ cortex. The average diameter at the main branch point of the apical dendrite was 3.1 μm, resembling measurements at electron microscopic resolution for layer 5 pyramidal neurons in mouse barrel cortex^46^. The total number of free parameters in each model was hence 35 (i.e., 32 from the conventional model reported by the Hay et al.,^13^, 2 for the decay of the Kv3.1 channel along the apical dendrite, and one for scaling its diameter).

### Simulations

Simulations of multi-compartmental models were performed with NEURON 7.4^47^ and custom written software in python. Temperature was set to 34°C, and the models where initialized with a membrane potential of −75 mV. The biophysical parameters (e.g. distributions of ion channels, intracellular calcium dynamic), and the diameters of the apical dendrite were optimized using the multi-objective evolutionary algorithm developed by Hay et al.,^13^ until simulated responses to current injections reproduced the characteristic responses of L5PTs within the empirically observed range **(Table S1)**. For optimization, the parameters were normalized with respect to the biophysically plausible ranges as reported by Hay et al.,^13^. For the 3 additional parameters, we used the following ranges: 0.5 to 3 for scaling of apical dendrite, 0 to 1 and −3 to 0 for the offset and slope of the Kv.3.1 channel density, respectively. Optimization was performed to capture the following dendritic physiology of L5PTs **(Fig. S2)**: backpropagating APs after a brief somatic current injection (1.9 nA, 5 ms), and Ca^2+^ APs during coincidence detection after a brief somatic current injection (1.9 nA, 5 ms) followed by an EPSP like current injection (5 ms delay, 0.5 nA) 180 μm below to the main branch point of the apical dendrite. These current injections were applied 295 ms after initialization of the simulation. To capture the characteristic perisomatic physiology of L5PTs, parameter optimization was additionally performed for somatic step current injections with the 0.619 nA, 0.793 nA and 1.507 nA amplitudes and a duration of 2 seconds. These current injections were applied 700 ms after initialization of the simulation. During the optimization, the 40 features of the simulated voltage traces were compared to the respective empirical data as proposed by Hay et al.,^13^, and as detailed in **Table S1**. The features were normalized by z-scoring with respect to the experimentally observed mean and standard deviation. Models were considered acceptable if the maximum z-scored deviation was less than 3.1 for the dendritic physiology, and less than 4.5 for the perisomatic physiology. The mean deviation of the thereby selected models was 1.05±0.14 **(Fig. S1)**, well below the cutoff criterion for acceptable models that was suggested previously^48^. We used the indicator based evolutionary (IBEA) algorithm^23^ for optimization with a population size of 1000, mutation probability of 0.3, and crossing-over probability of 0.7. We used dask to parallelize the optimization. The optimization was performed in two steps. First, optimization was performed for 600 generations to capture the dendritic physiology. Second, if models with a maximum deviation of <3 standard deviations were found, we continued the optimization by incorporating the perisomatic physiology into the objectives. The optimization was terminated if there was no progress, or when acceptable models were found. Repeating the optimization procedure with different seeds generally resulted in new models, which extended the parameter ranges for each morphology. We continued this parameter space exploration until the parameter range coverage was close to 100% (for models constrained for dendritic and perisomatic physiology), and the principal component space was densely filled (for models constrained for dendritic physiology). In total, 15,569,645 were generated that were constrained for dendritic physiology, of which 4,239,392 in addition were constrained for perisomatic physiology.

### Qantification and statistical analysis

For all models with acceptable dendritic physiology (i.e., including those with acceptable perisomatic physiology), we extracted the membrane potential, currents through active ion channels, and the intracellular calcium concentration at the soma and main branch point of the apical dendrite during simulations of coincidence detection. From these data, we computed the total charge exchanged (unit: Coulomb) through each of the seven ion channels that are expressed at the main branch point by integrating the currents for a 50 ms time interval starting with the onset of the somatic current injection. Currents through Ih channels were negligible in comparison to the other six ionic currents, and discarded from further analysis (the density Ih channels was also not subject to optimization). The charge exchanged through the remaining six ion channels was then used to compute the first two principal components across all acceptable models. Among the three depolarizing channels – voltage-gated sodium channel, high- and low-voltage activated calcium channels – the latter were dominating the generation of Ca^2+^ APs during coincidence detection across all models. Thus, we computed the depolarization index in **Fig. 2c** only for the calcium channels:

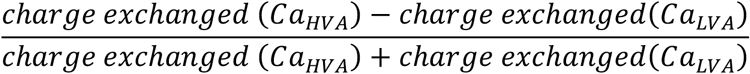

Similarly, M and SK channels dominated hyperpolarization across the majority of models, and we hence computed the hyperpolarization index in **Fig. 2d** only for these two potassium channels. To estimate the energy cost of dendritic Ca^2+^ APs, we computed the difference between membrane potential and the reversal potential of each ionic species (potassium: −85 mV, sodium: 50 mV, calcium: the reversal potential was constantly updated based on the varying intracellular Ca^2+^ concentration using the Nernst equation). These potential differences were multiplied with the respective ionic currents and integrated over the 50 ms time interval using Simpson’s rule. Correlations and associated p-values were computed using the method “pearsonr” of the scipy python package (version 0.18.2).

## Acknowledgments

We thank Bert Sakmann for comments on the manuscript, Etay Hay for providing the conventional biophysical model and genetic algorithm, and Christiaan de Kock for providing some of the morphologies. Funding was provided by the Center of Advanced European Studies and Research, the German Federal Ministry of Education and Research (grant 01IS18052, to M.O.), the Deutsche Forschungsgemeinschaft (SFB 1089 and SPP 2041, to M.O.), and the European Research Council under the European Union’s Horizon 2020 research and innovation program (grant 633428, to M.O.).

## Author contributions

M.O. and A.B. conceived and designed the study. A.B. developed the model, performed simulations, and analyzed the data. M.O. and A.B. wrote the paper.

The authors declare no competing interests.

---

**Fig. S1.**
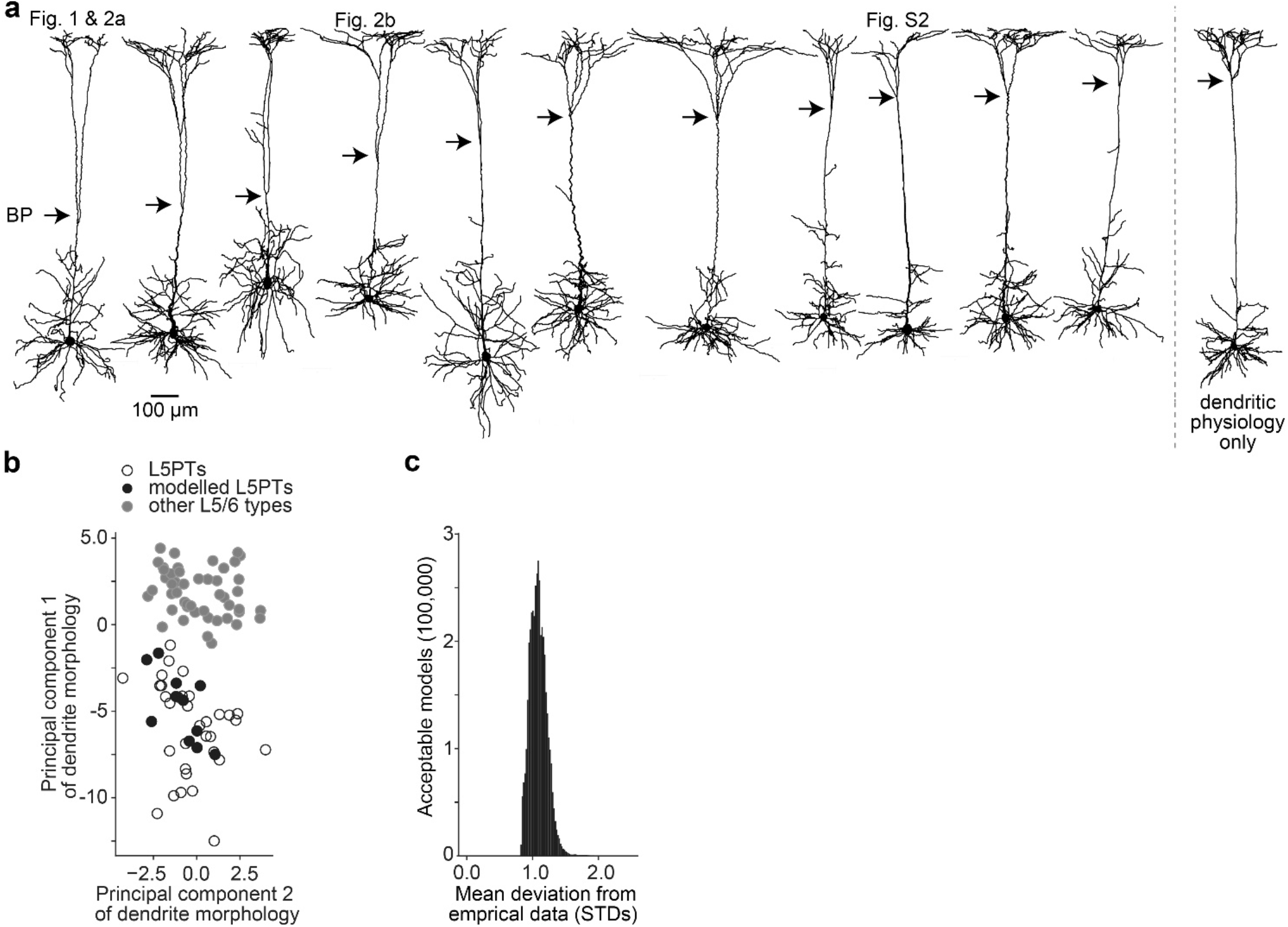
(related to Fig. 1): Models represent the morphological diversity of L5PTs. **a.** Gallery of the *in vivo*-labeled dendrite morphologies used in this study. The main branch point (BP) of the apical dendrite reflects the initiation zone for Ca^2+^ APs. For one of these morphologies (most right), we could generate models that reproduce the dendritic physiology of L5PTs, but no models that reproduce both dendritic and perisomatic physiology. **b.** The morphologies for which we generated models represent the morphological diversity of L5PTs in the rat barrel cortex. Here we visualize the first two principal components derived from 22 morphological features that reliably discriminate L5PTs (n=44) from the other major excitatory cell types in layers 5/6 (n=47; i.e., L5ITs, L6CCs, L6CTs)^26^. **c.** Mean error across all models with acceptable dendritic and perisomatic physiology, quantified as the deviation from empirical mean in units of standard deviations across all 40 objectives **(Table S2)**.

**Fig. S2.**
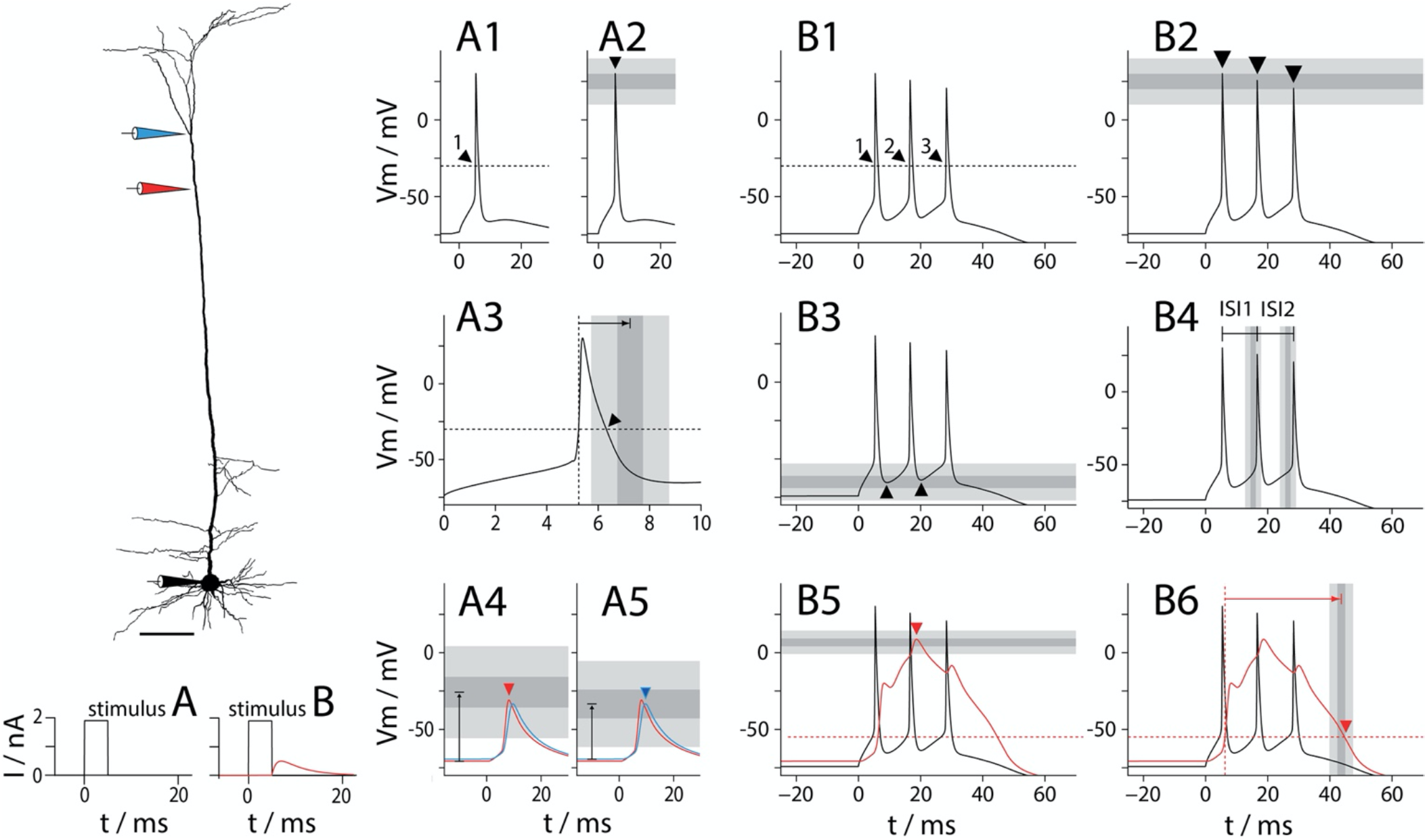
(related to Fig. 2): Empirical constraints for models with acceptable dendritic physiology. Panels A1-B6: features extracted from example simulations of responses to stimulus A (5 ms somatic step current injection of 1.9 nA) and stimulus B (as stimulus A, but combined with an EPSP like injection of 0.5 nA into the Ca^2+^ zone of the apical dendrite). Grey corridors represent the mean of the empirical data ± one standard deviation (dark grey), and three standard deviations (light grey) **A1.** Stimulus A evokes a single AP. **A2.** AP peak is 25±5 mV. **A3.** AP width is 2±0.5 ms. **A4.** Amplitude of backpropagating AP at the red recording pipette (located 180 μm below the main branch point) is 45±10 mV. **A5.** Amplitude at the blue recording pipette (located at the main branch point) is 36±9.33 mV. **B1.** Stimulus B evokes a burst of three axonal APs. **B2.** The mean AP peak across all 3 APs is 25±5 mV. **B3.** Mean somatic after-hyperpolarization (AHP) depth is −65±4 mV. **B4.** The mean somatic inter-spike-interval (ISI) is 9.9±0.85 ms. **B5.** The Ca^2+^ AP peak is 6.73±2.54 mV. **B6.** The Ca^2+^ AP width is 37.43±1.27 ms.

**Fig. S3.**
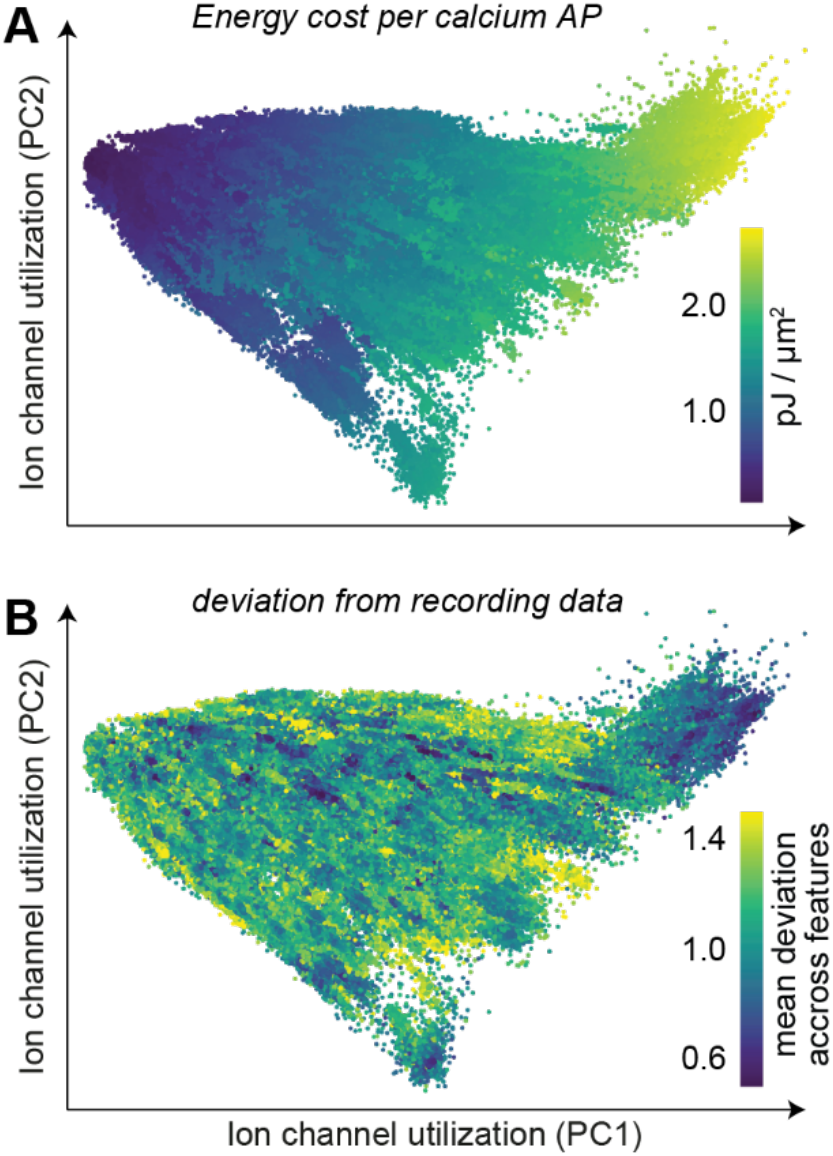
(related to Fig. 3): Energy-efficient and inefficient implementations are equally realistic in their response properties. **a.** Principle component space as in **Fig. 2** and **Fig. 3**. Energy efficiency correlates with the first principal component. **b.** There is no correlation between the “realism” of a model’s response as quantified by the mean deviation of its responses from experimental traces and its position in the principal component space.

**Fig. S4.**
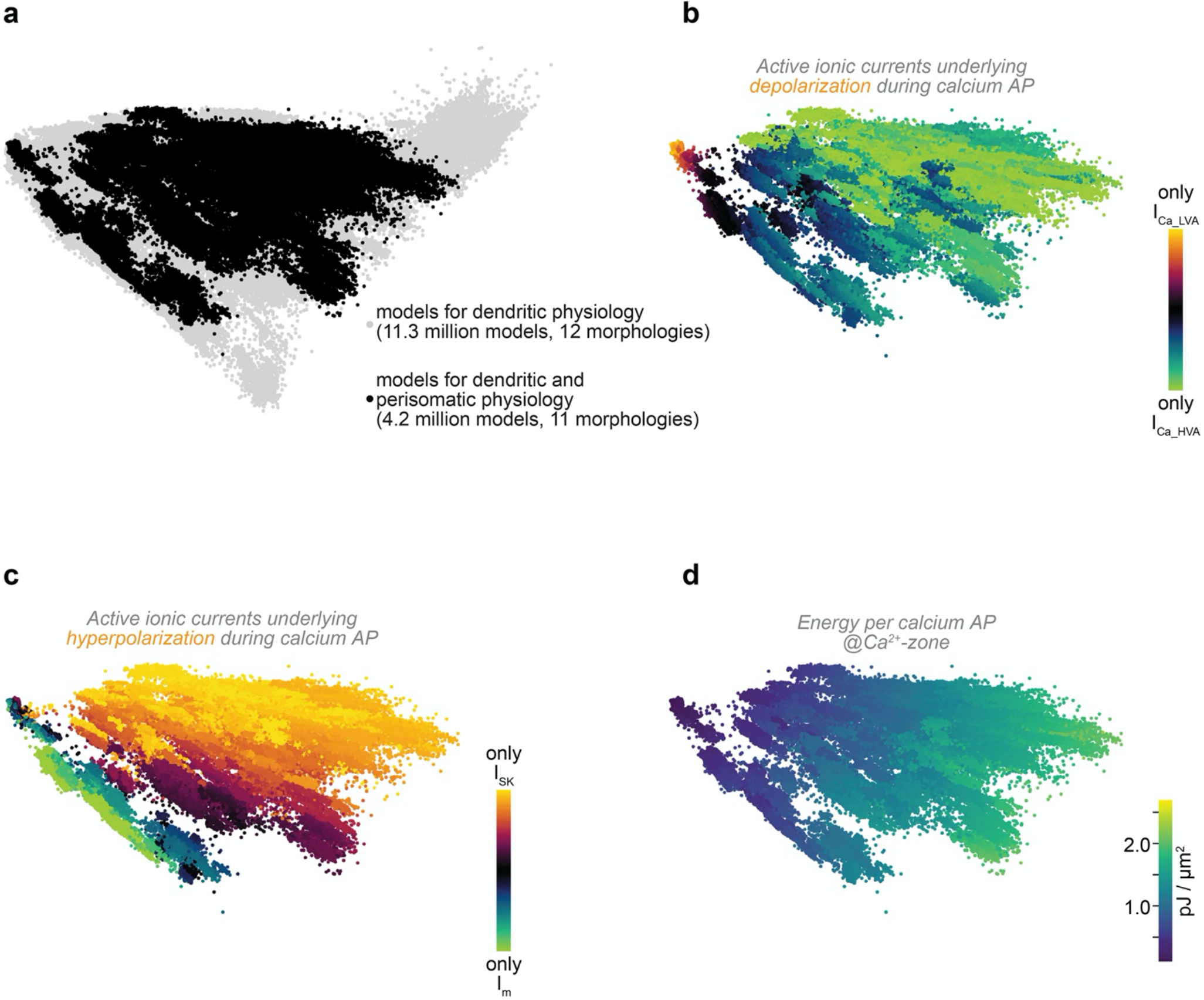
(related to Fig. 3): Principles underlying the dendritic physiology of L5PTs are independent from the perisomatic physiology. **a.** Principle component space as in **Fig. 2** and **Fig. 3**. Models which reproduce only the dendritic physiology of L5PTs (n=11.3 million, grey dots) share the same space and intermingle with those models that capture both the dendritic and perisomatic physiology (n=4.2 million, black dots). This indicates that perisomatic properties do not constrain the channel utilization required to capture the dendritic physiology. **b-d.** Same analysis as in **Fig. 2** and **Fig. 3**, but restricted to the 4.2 million models that reproduce both the dendritic and perisomatic physiology. Consistent with **Fig. 2c**, models transition along principal component 1 from Ca_LVA as the predominant depolarizing mechanism to Ca_HVA (panel b). Consistent with **Fig. 2d**, models transition along principal component 2 from M currents as the dominating hyperpolarizing mechanism to calcium-dependent potassium currents (panel c). Consistent with **Fig. 3a**, the energy cost per Ca^2+^ AP increases along principal component 1 (panel d). Thus, the organizational principles underlying the generation of Ca^2+^ APs are largely unaffected by constraints that are required to capture the perisomatic physiology.

**Table S1:** Acceptable models match the empirically observed mean and standard deviation across features of dendritic and perisomatic physiology. Empirical data (grey) was adopted from Hay et al. 2011^13^. Please note: matching the means and standard deviations across models with those observed empirically across L5PTs was not an objective during the parameter space exploration. Remarkably, the distributions of parameters across all acceptable models resemble nonetheless those observed empirically across L5PTs (i.e., similar means and standard deviations for most features).

| Features of perisomatic physiology |  |  |  | Features of dendritic physiology |  |
| --- | --- | --- | --- | --- | --- |
| Feature | Mean ± STD | | | Feature | Mean ± STD |
|  | current step 1 | current step 2 | current step 3 |  |  |
| 1. AP frequency (Hz) | 9 ± 0.88 | 14.5 ± 0.56 | 22.5 ± 2.22 | 1. Ca<sup>2+</sup> AP peak (mV) | 6.73 ± 2.54 |
| | 9.4 ± 0.83 | 14.3 ± 0.72 | 24.6 ± 1.83 | | 7.36 ± 1.92 |
| 2. Adaptation index | 0.0036 ± 0.0091 | 0.0023 ± 0.0056 | 0.0046 ± 0.0026 | 2. Ca<sup>2+</sup> AP width (mV) | 37.43 ± 1.27 |
| | 0.0078 ± 0.0088 | 0.0071 ± 0.0053 | 0.0086 ± 0.0035 | | 37.63 ± 0.96 |
| 3. ISI-CV | 0.1204 ± 0.0321 | 0.1083 ± 0.0368 | 0.0954 ± 0.0140 | 3. Somatic AP spike count | 3 ± 0 |
| | 0.0866 ± 0.0369 | 0.0936 ± 0.0281 | 0.1165 ± 0.0198 | (coincident current injection) | 3 ± 0 |
| 4. Initial Burst ISI (ms) | 57.75 ± 33.48 | 6.625 ± 8.65 | 5.38 ± 0.83 | 4. Somatic AP ISI (ms) | 9.9 ± 0.85 |
| | 43.66 ± 14.41 | 17.17 ± 5.62 | 6.56 ± 0.72 | | 10.1 ± 10.7 |
| 5. First spike latency (ms) | 43.25 ± 7.32 | 19.13 ± 7.31 | 7.25 ± 1.00 | 5. Somatic AHP depth (mV) | -65 ± 4 |
| | 34.38 ± 5.41 | 20.26 ± 2.51 | 7.01 ± 0.89 | | -65 ± 1 |
| 6. AP peak (mV) | 26.23 ± 4.97 | 16.52 ± 6.11 | 16.44 ± 6.93 | 6. Somatic AP peak (mV) | 25 ± 5 |
| | 20.77 ± 3.50 | 20.58 ± 3.56 | 19.23 ± 3.85 | | 27 ± 3 |
| 7. Fast AHP depth (mV) | -51.95 ± 5.82 | -54.19 ± 5.57 | -56.56 ± 3.58 | 7. Somatic AP half-width (ms) | 2 ± 0.5 |
| | -61.68 ± 1.17 | -61.53 ± 1.21 | -60.52 ± 1.43 | | 1 ± 0.1 |
| 8. Slow AHP depth (mV) | -58.04 ± 4.58 | -60.51 ± 4.67 | -59.99 ± 3.92 | 8. Somatic AP spike count | 1 ± 0 |
| | -61.96 ± 1.23 | -61.63 ± 1.28 | -60.38 ± 1.43 | | 1 ± 0 |
| 9. Slow AHP time | 0.238 ± 0.030 | 0.279 ± 0.027 | 0.213 ± 0.037 | 9. bAP amplitude at 620 μm (mV) | 45 ± 10 |
| | 0.225 ± 0.039 | 0.258 ± 0.037 | 0.215 ± 0.054 | 180 μm below the BP | 33 ± 8 |
| 10. AP half-width (ms) | 1.31 ± 0.17 | 1.38 ± 0.28 | 1.86 ± 0.41 | 10. bAP amplitude at 800 μm (mV) | 36 ± 9.33 |
| | 0.96 ± 0.04 | 0.96 ± 0.04 | 0.96 ± 0.04 | at the BP | 23 ± 7.72 |

**Table S2:**
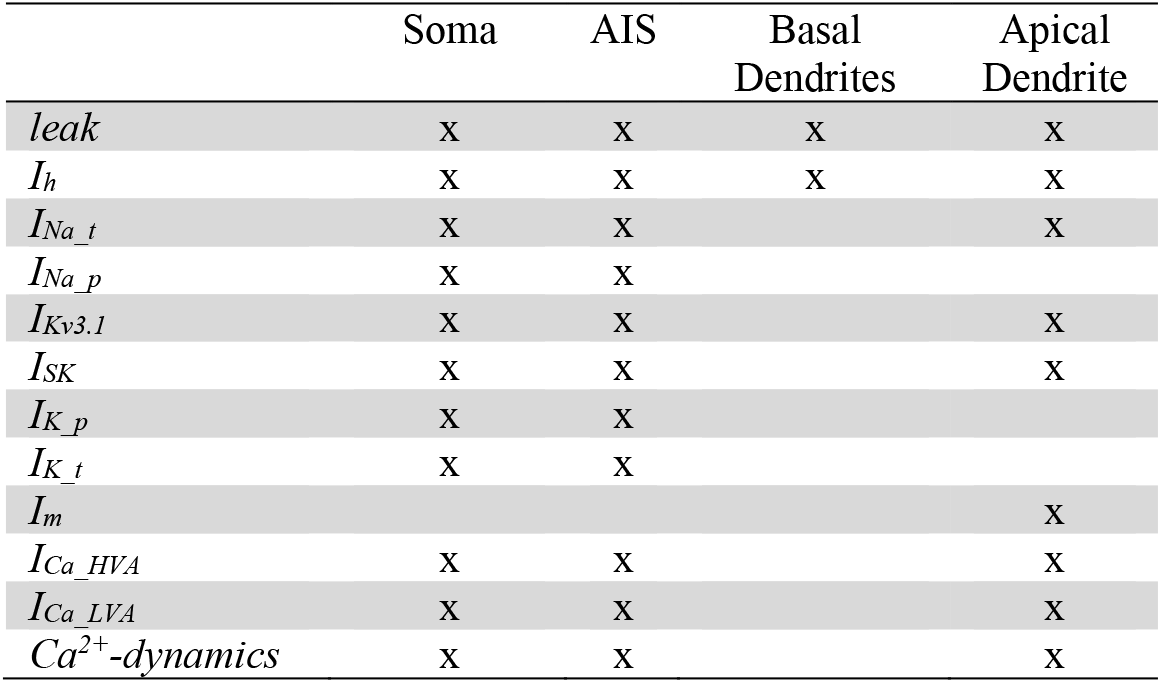
Ionic currents by subcellular compartment. Abbreviations: Axon initial segment, AIS. Non-specific cation current, I_h_. Fast inactivating Na^+^ current, I_Na_t_. Persistent Na^+^ current, I_Na_p_. Fast non-inactivating K^+^ current, I_Kv3.1_. Small-conductance Ca^2+^ activated K^+^ current, I_SK_. Slow inactivating K^+^ current, I_K_p_. Fast inactivating K^+^ current, I_K_t_. High-voltage activated Ca^2+^ current, I_Ca_HVA_. Low-voltage activated Ca^2+^ current, I_Ca_LVA_. Muscarinic K^+^ current, I_m_. Intracellular [Ca^2+^] dynamics.

|  | Soma | AIS | Basal Dendrites | Apical Dendrite |
| --- | --- | --- | --- | --- |
| *leak* | x | x | x | x |
| *I<sub>h</sub>* | x | x | x | x |
| *I<sub>Na_t</sub>* | x | x |  | x |
| *I<sub>Na_p</sub>* | x | x |  |  |
| *I<sub>Kv3.1</sub>* | x | x |  | x |
| *I<sub>SK</sub>* | x | x |  | x |
| *I<sub>K_p</sub>* | x | x |  |  |
| *I<sub>K_t</sub>* | x | x |  |  |
| *I<sub>m</sub>* |  |  |  | x |
| *I<sub>Ca_HVA</sub>* | x | x |  | x |
| *I<sub>Ca_LVA</sub>* | x | x |  | x |
| *Ca<sup>2+</sup>-dynamics* | x | x |  | x |

## Notes

### Competing Interest Statement

The authors have declared no competing interest.

